# Nuclei isolation protocol from diverse angiosperm species

**DOI:** 10.1101/2022.11.03.515090

**Authors:** Chenxin Li, Joshua C. Wood, Natalie C. Deans, Anne Frances Jarrell, Dionne Martin, Kathrine Mailloux, Yi-Wen Wang, C. Robin Buell

**Affiliations:** Center for Applied Genetic Technologies, University of Georgia, Athens GA, 30602; Department of Crop and Soil Sciences, University of Georgia, Athens GA, 30602

**Keywords:** plant nuclei isolation, single cell genomics

## Abstract

The ability to generate intact nuclei is crucial to the success of a variety of genomics experiments, such as Assay for Transposase-Accessible Chromatin using sequencing (ATAC-seq), Cleavage Under Targets and Tagmentation (CUT&Tag), and nuclei-based single cell sequencing (e.g., single nuclei ATAC-seq and single nuclei RNA-seq). For plants, the presence of the cell wall presents significant challenges in the isolation of nuclei from tissues. Here, we report an optimized nuclei isolation protocol that can be adapted for diverse angiosperm species, including maize, soybean, tomato, potato, and wheat, starting from fresh or frozen tissues. Nuclei release is achieved through chopping tissue on ice, where a key parameter affecting nuclei integrity is the concentration of detergent TritonX-100 in the nuclei isolation buffer. The method is simple, quick, and largely centrifugation-free, in which debris is removed by serial filtration. Initial nuclei release and filtration can be performed within 20 min. Fluorescence activated nuclei sorting is then used for final nuclei purification to remove other organelles such as plastids. The protocol uses 500 mg or less plant tissue as input and typically yields at least 100,000 – 200,000 purified nuclei per sample, a common input amount for downstream experiments. Throughout the protocol, we provide guidelines for optimization if performing nuclei isolation from a given species and tissue for the first time.

## Introduction

Recently, there have been rapid developments in the techniques for studying epigenomics and single cell genomics (Huang and Ecker 2018; Shema, Bernstein, and Buenrostro 2019; Luo, Fernie, and Yan 2020; Preissl, Gaulton, and Ren 2022). Multiple techniques have been developed to investigate gene expression, chromatin accessibility, chromatin modifications, and transcription factor binding, such as Assay for Transposase-Accessible Chromatin using sequencing (ATAC-seq) (Grandi et al. 2022), Isolation of Nuclei Tagged in Specific Cell Types (INTACT) (Deal and Henikoff 2010), Cleavage Under Targets and Release Using Nuclease (CUT&RUN) (Meers et al. 2019), Cleavage Under Targets and Tagmentation (CUT&Tag) (Kaya-Okur et al. 2019; Ouyang et al. 2022), and single nuclei RNA-seq/ATAC-seq (Grindberg et al. 2013; Kashima et al. 2020; Buenrostro et al. 2015). A common input for these experiments is purified nuclei from unfixed, non-crosslinked tissues. Intact nuclei are crucial to the success of these techniques, as damaged nuclei may lead to RNA leakage and deterioration of chromatin conformation, and as a consequence, sub-optimal results. The isolation of intact nuclei is relatively routine from animal soft tissues and cultured cells. However, for plants, the presence of cell walls impedes the isolation of nuclei, since a harsher tissue disruption method such as grinding, agitation, or chopping, is required to break open cell walls. In addition, the presence of plastids, lipid bodies, and starch granules within plant cells may also complicate nuclei isolation.

We attempted nuclei isolation with several published nuclei isolation protocols (Marand et al. 2021; Thibivilliers, Anderson, and Libault 2020; Sikorskaite et al. 2013) yet did not recover robust nuclei with our target species. During troubleshooting, we observed that the tissue disruption method, the concentration of detergent, as well as centrifugation and resuspension steps significantly influenced the integrity of isolated nuclei. Here, we present a modified and benchmarked nuclei isolation protocol that was tested across a range of angiosperm species. The protocol (Figure 1) uses chopping as the means to release nuclei from tissues, followed by a serial filtration to remove debris. The protocol is quick. The initial tissue disruption and filtration steps can be performed within 20 min. The resultant crude nuclei suspension is then subjected to fluorescence activated nuclei sorting for further nuclei purification. From 500 mg or less of input material, the protocol typically yields at least 100,000 – 200,000 nuclei from 1 mL of lysate. The critical parameter affecting nuclei integrity is the detergent (TritonX-100) concentration. Insufficient detergent concentration leads to under-lysis of cells in which nuclei are still enclosed in cells, while excessive concentrations of detergent lead to over-lysis and damaged nuclei. We have tested and optimized this protocol for a collection of diverse angiosperm species, including major crops. Optimal TritonX-100 concentrations and demonstration of nuclei integrity are reported below.

**Figure 1:**
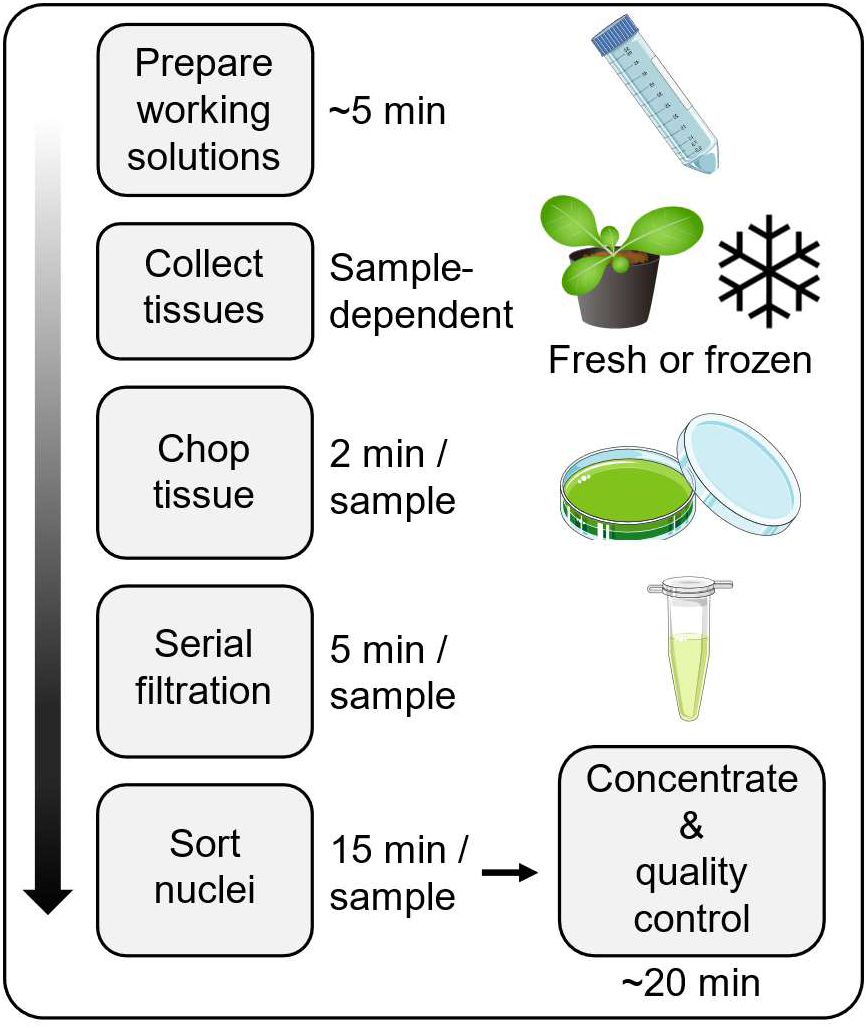
Overview and timeline of the protocol. This figure was prepared using stamps available online at the Biocon repository (https://bioicons.com/).

## Materials and Methods

### Reagents

- Tris-HCl solution, 1 M, pH 8 (Invitrogen 15568025)
- EDTA disodium salt dihydrate (Sigma-Aldrich, E5134)
- KCl (Sigma-Aldrich, P9541)
- NaCl (Sigma-Aldrich, S9888)
- Spermine tetrachloride (Sigma-Aldrich, S2876)
- TritonX-100 (Sigma-Aldrich, X100)
- 2-mercaptoethanol (2-ME, Sigma-Aldrich, M6250)
- 4’,6-Diamidino-2-phenylindole dihydrochloride (DAPI, Sigma-Aldrich, D9542)
- Bovine Serum Albumin (BSA, Sigma-Aldrich, A9430)

### Stock solutions

- EDTA, 0.5 M, pH 8
- KCl, 1 M
- NaCl, 1M
- TritonX-100, 10% v/v
- Spermine tetrachloride, 1 M
- DAPI, 1 mg/mL (1000X concentrate)

### Consumables

- 1.5 mL microcentrifuge tubes
- 50 mL conical tubes
- 100 μm cell strainer (Fisher Scientific 22-363-549)
- 40 μm cell strainer (Fisher Scientific 22-363-547)
- 20 μm cell strainer (pluriSelect 43-50020-03)
- 1000 μL and 200 μL wide bore pipette tips
- Serological pipettes and controller
- Petri dishes
- Razor blades
- Microscope slides and cover slips

### Working solutions, keep on ice

The following recipes were modified from Marand et al. (2021).

### Methods

1. Place 0.5 g of fresh or frozen tissue on a petri dish on ice*. * More input does not always lead to higher yield. Keep samples on ice as much as possible. Once the tissues are harvested and placed on ice, process them quickly, ideally < 30 min.
2. Transfer 1000 μL NIB to petri dish. Place tissue into NIB in petri dish. Use a pair of forceps to handle and wet the tissue with NIB (Figure 2A, C).
3. Use a single edge razor blade to rapidly chop the tissue with force for 2 min. The optimal chopping frequency is ~200 – 250 beats per minute (bpm). By the end of the 2 min period, the tissue will turn into a fine slurry resembling the texture of pesto (Figure 2B, D). * Insufficient tissue disruption can lead to low nuclei yield; however, we strongly discourage chopping for more than 2 min, which can cause mechanical damage to nuclei.
4. Add 1000 μL NIB to petri dish and gently rinse minced tissue.
5. Wet 100 μm, 40 μm, and 20 μm strainers with 2 mL NIB.
6. Pass the nuclei slurry through the 100 μm strainer*. * Always use wide bore tips to transfer nuclei. Pipette slowly.
7. Slowly pass the 100 μm filtrate through the 40 μm strainer using a serological pipette.
8. Slowly pass the 40 μm filtrate through the 20 μm strainer using a serological pipette.
9. Rinse the 20 μm strainer with deionized water and tap clean on a paper towel.
10. Slowly pass the 20 μm filtrate through the 20 μm strainer again.
11. Transfer 1 mL filtrate to a new 1.5 mL tube and stain with 1 μL 1 mg/μL DAPI*. * 1 mL of filtrate will typically yield at least 100,000 nuclei from 0.5 g of tissue. Adjust DAPI volume (1000x concentrate) if using a different volume of filtrate in downstream experiments.
12. Recommended if performing nuclei from a given species and tissue for the first time or optimizing TritonX-100 concentrations: Check nuclei integrity under a fluorescence microscope with DAPI fluorescence (40× objective lens or higher magnification). Intact nuclei should have clear edges with minimal blebbing or chromatin extrusion (Figure 3). Occasionally, weakly DAPI-stained regions representing nucleolus (Kodiha, Bański, and Stochaj 2011) and strongly DAPI-stained spots representing heterochromatin (Probst et al. 2003) are visible (Figure 3).
13. Nuclei purification using flow cytometry: We use a Moflo Astrios EQ running preservative-free flow cytometry solution. The following gating strategy and parameters were modified from Marand et al. (2021).
  a. Transfer 500 μL NSB into a new 1.5 mL tube, which will be used as the catch tube*. * Insufficient volume of liquid in catch tube may lead to mechanical damage when nuclei land in the catch tube. Alternatively, a reaction buffer of the following experimental step may be used in place of NSB, such as 10x Genomics Nuclei Buffer (10x Genomics Chromium Single Cell Multiome ATAC + Gene Expression Kit).
  b. The instrument was set up with a 100 μm tip and sheath pressure at 25 psi.
  c. Nuclei were run below 500 events per second (EPS) depending on the concentration*. * High EPS may lead to mechanical damage to nuclei.
  d. Nuclei were gated on log scaled forward scatter (FSC) vs. log scaled side scatter (SSC) to eliminate debris (Figure 4A).
  e. DAPI width vs. DAPI area gating was used to reduce potential doublets (Figure 4B).
  f. Green fluorescence vs. DAPI area gating was used to reduce green fluorescent material (Figure 4C).
  g. The final gate was on DAPI area selecting the first strong peak and all subsequent peaks corresponding to different cell cycle or endoreduplication stages (Figure 4D).
  h. Sorting was performed in Purity 1 drop mode and triggered on DAPI fluorescence.
  i. We aim for between 100,000 to 200,000 nuclei per sample.
14. Concentrate nuclei by a final centrifugation step.

a. Centrifuge nuclei in catch tubes at 300 × g for 10 min in a refrigerated centrifuge at 4°C*. * Repeated centrifugation and resuspension can cause mechanical damage to nuclei. Thus, we suggest minimal centrifugation throughout the protocol.
b. Carefully remove supernatant with a wide bore tip*. The pellet may be invisible. * Based on the desired final concentration, a small amount of NSB is retained. For example, if 200,000 nuclei were sorted and the target concentration is > 5000 nuclei/μL, 40 μL or less NSB should be retained in the tube.
c. Resuspend nuclei by gently tapping the tube.
d. Check nuclei integrity under a microscope using DAPI fluorescence (40×) objective lens or higher magnification).

**Figure 2:**
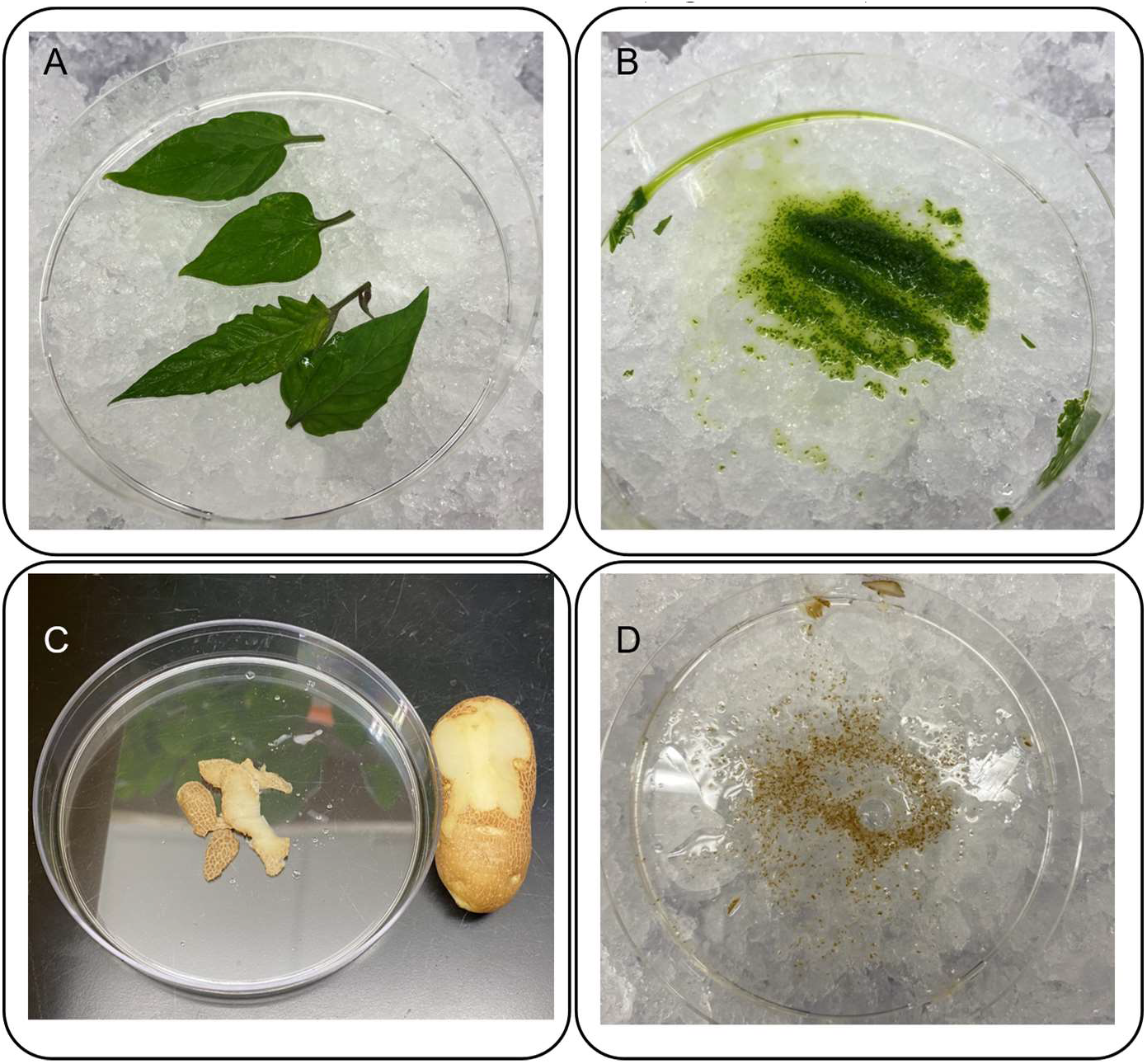
Tissue disruption via chopping. (A) Tomato leaflets on a petri dish. (B) The same tomato sample in (**A**) after 2 min of chopping in NIB. (C) Peels from a potato tuber. (D) The same potato sample in (**C**) after 2 min of chopping in NIB.

**Figure 3:**
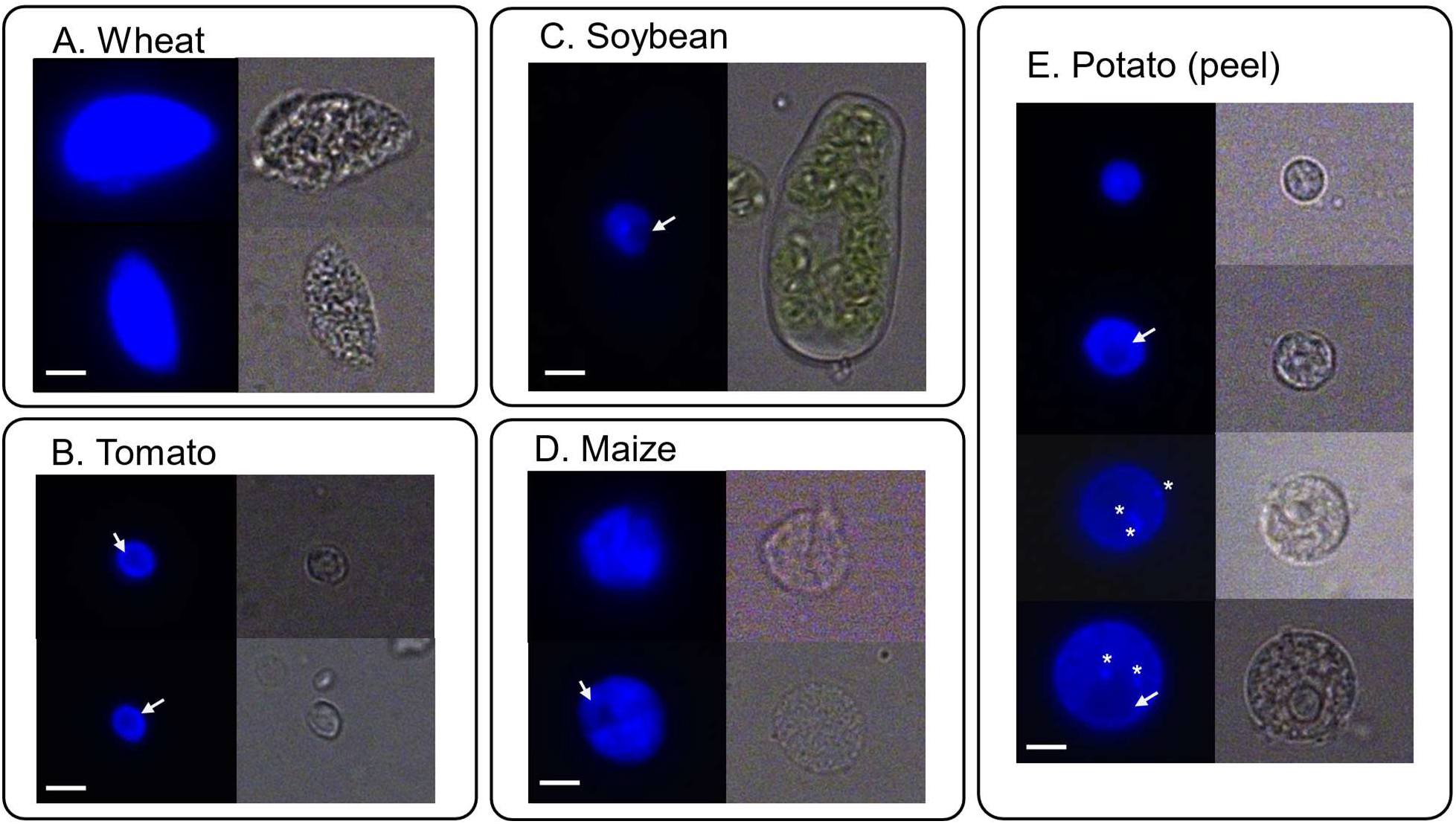
Representative images of intact nuclei after serial filtration and DAPI staining. (A) Nuclei from seedling leaves of hexaploid wheat. (B) Nuclei isolated from tomato leaflets. (C) A nucleus trapped in an under-lysed cell from soybean seedling leaves. (D) Nuclei from maize seedling leaves. (E) Different size nuclei isolated from potato tuber peels, potentially corresponding to different endoreduplication states. Left images: blue signal: DAPI fluorescence. Right images: Bright field. Bars: 5 μm. Arrows: regions in the nuclei with weak DAPI signal, likely nucleolus. Asterisks: bright spots in the nuclei with strong DAPI signal, likely heterochromatic regions.

**Figure 4.**
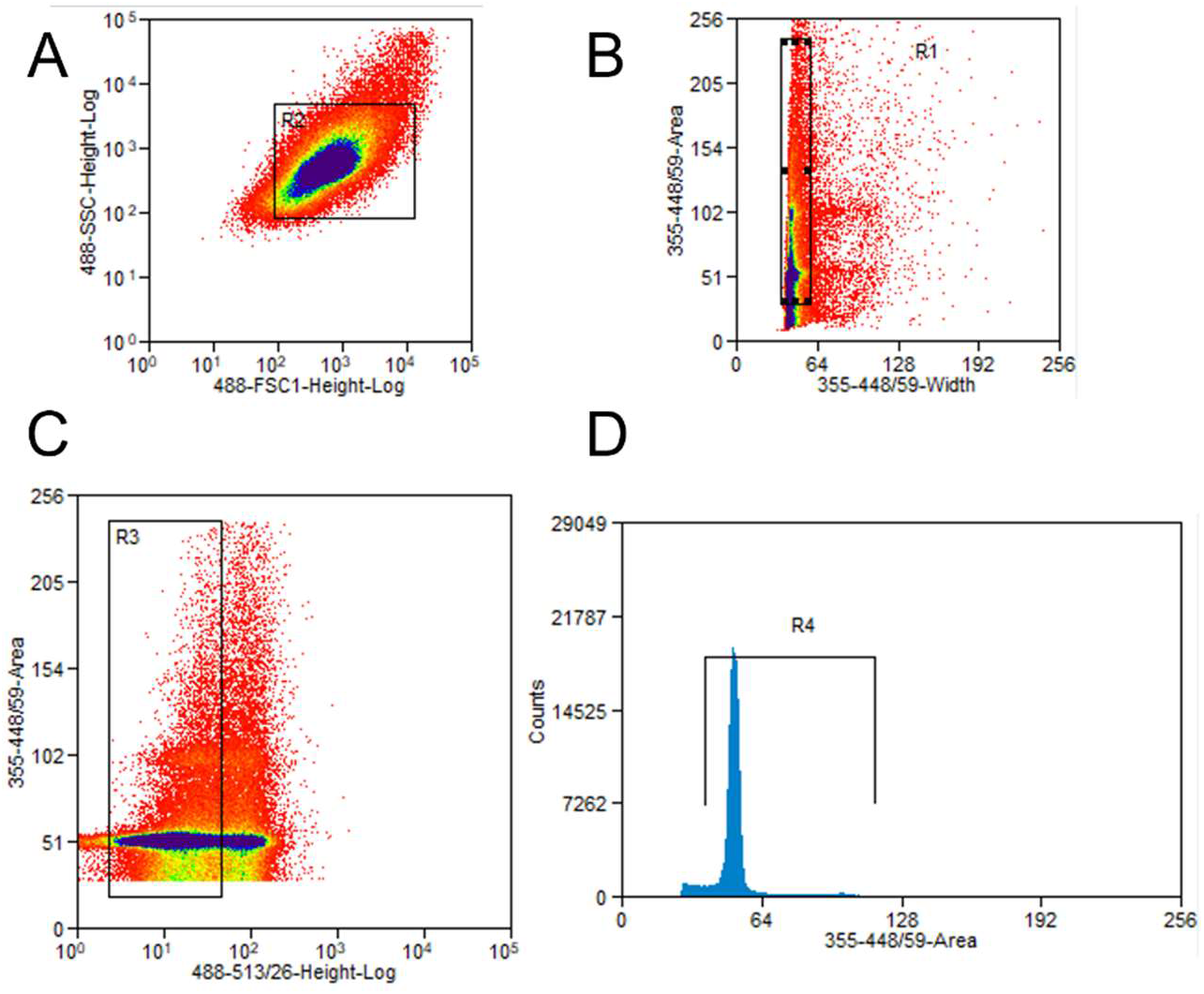
Purification of nuclei from organelles using fluorescence activated sorting. (A) Forward scatter vs. side scatter eliminates debris. (B) DAPI width vs. DAPI area removes aggregated nuclei. (C) Green fluorescence vs DAPI area to remove green fluorescent material. (D) Distribution of DAPI area. Regions on graphs included by rectangle (A-C) or bar (D) indicate selected droplets. Droplets outside of selected regions are discarded.

## Results and Discussion

We have determined the optimal TritonX-100 concentration across a range of angiosperm species spanning five taxonomic families. Overall, there was no association between either estimated genome size or taxonomic group and optimal detergent concentrations (Table 3). However, we observed that species with larger estimated genome size also had larger nuclei (Figure 3). In addition, we speculate that different nuclei sizes from the same sample may reflect different endoreduplication states (Sugimoto-Shirasu and Roberts 2003), as shown in potato peel samples (Figure 3E). We examined nuclei integrity under a 100 × objective lens from at least two independent replicates. Overall, we observed at least 70% intact nuclei from a random sample of 10 – 17 nuclei per replicate (Figure 5). The exact percentages of intact nuclei may differ depending on the individual who performs the experiment, the hydration status of the sample, the age and health of the plants, and the amount of input, among other factors. In our experience, different individuals yielded very similar outcomes if they used the species/tissue-optimized TritonX-100 concentration. The protocol appeared to perform well for both above ground (leaf and leaflet) and below ground tissues (potato peel) (Figure 5); however, the protocol has not been optimized for woody tissues such as stems and roots with pronounced secondary growth. For the soybean leaf sample specifically, we observed under-lysed cells in which nuclei were trapped (Figure 3C). In our hands, increasing the detergent concentration resulted in more lysed free nuclei, while reducing detergent concentration resulted in more under-lysed cells.

**Figure 5.**
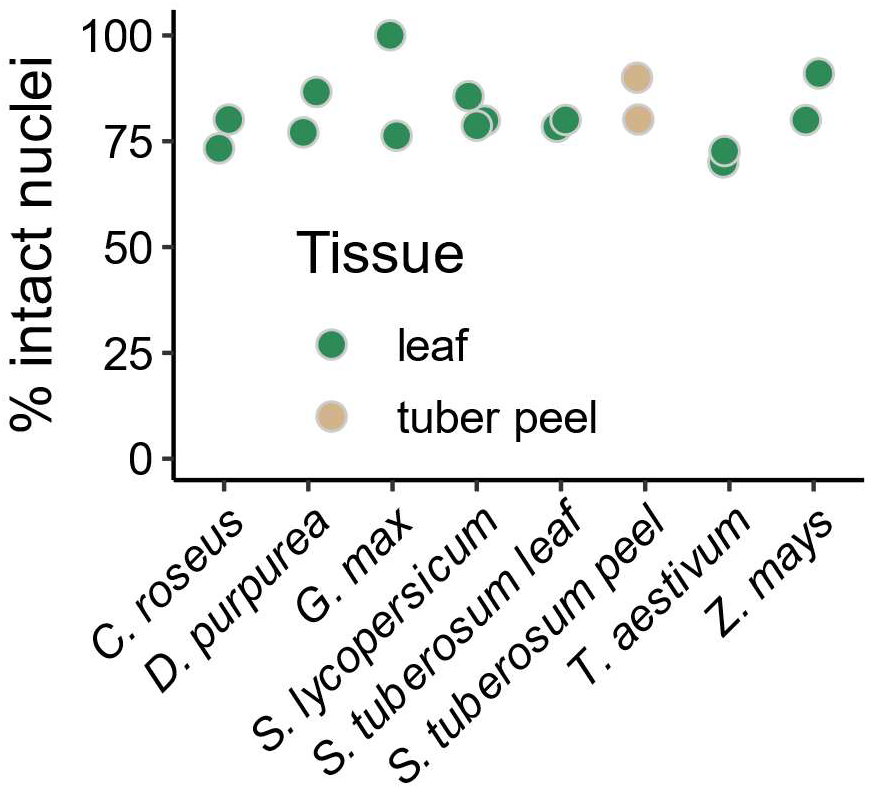
Percentage of intact nuclei (as exemplified in Figure 3) across materials processed by this protocol, using optimal TritonX-100 listed on Table 3. Each data point is a random sample of 10 - 17 nuclei from an independent biological replicate.

**Table 1:** Nuclei Isolation Buffer (NIB)

| Component | Volume (for 25m L) | Final Concentration |
| --- | --- | --- |
| Tris-HCl, 1 M, pH 8 | 375 µL | 15 mM |
| EDTA 0.5 M, pH 8 | 100 µL | 2 mM |
| KCl, 1 M | 2 mL | 80 mM |
| NaCl, 1 M | 0.5 mL | 20 mM |
| 2-ME (100%) | 10 µL | 0.04% v/v |
| Spermine, 1M | 12.5 µL | 0.5 mM |
| BSA | 25 mg | 0.1% w/v |
| TritonX-100, 10% | Variable* | 0.01% to 0.15% v/v |
| H2O | Fill up to 25 mL |  |

\* The optimal TritonX-100 concentration varies between species and tissues. If isolating nuclei from a given species and tissue for the first time, we suggest testing a series of TritonX-100 concentrations (e.g., 0.01% to 0.15%) to determine the optimal concentration.

**Table 2:**
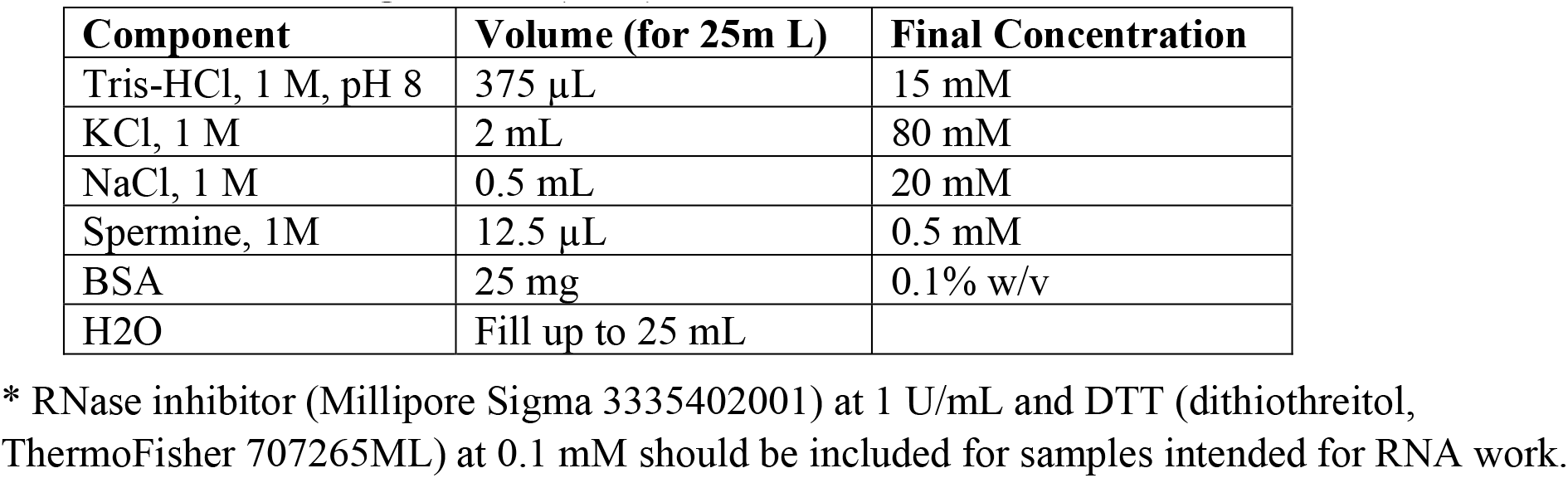
Nuclei Storage Buffer (NSB)*

**Table 3.** Metadata and optimal TritonX-100 concentrations of materials tested. Estimated genome size from (Guimarães et al. 2012; Castro and Castro 2012; Shultz 2006; Barone et al. 2008; Hoopes et al. 2022; Haberer et al. 2005; Zimin et al. 2017).

| Species | Tissue | Stage | Est. Genome Size (1c) | Optimal TritonX-100 Conc. | Family |
| --- | --- | --- | --- | --- | --- |
| *Catharanthus roseus* (Madagascar Periwinkle) | Leaf | 6-week | 738 Mb | 0.01% | Apocynaceae |
| *Digitalis purpurea* (foxglove) | Leaf (frozen) | Mature | 940 Mb | 0.0375% | Plantaginaceae |
| *Glycine max* (soybean) | Leaf | 2-week | 1.1 Gb | 0.1% | Fabaceae |
| *Solanum lycopersicum* (tomato) | Leaflet | 2-week | 950 Mb | 0.0375% | Solanaceae |
| *Solanum tuberosum* (potato) | Leaflet | 4-week | 4n = 3.2 Gb | 0.0375% | Solanaceae |
| *Solanum tuberosum* (potato) | Tuber peel | Developing tuber | 4n = 3.2 Gb | 0.0375% | Solanaceae |
| *Triticum aestivum* (wheat) | Leaf | 2-week | 17 Gb | 0.0375% | Poaceae |
| *Zea mays* (maize) | Leaf | 2-week | 2.4 Gb | 0.75% | Poaceae |

### Appendix: growth conditions of plants used in work

*Catharanthus roseus* plants were grown in a growth chamber under 24°C constant temperature, 15-hr day, and 300 μM light intensity. *Digitalis purpurea* plants were grown in a greenhouse with 24°C daytime temperature, 16°C nighttime temperature, 15-hr days, and 600 μM light intensity. Soybean, potato, wheat, and maize plants were grown in a growth chamber under 25.5°C daytime temperature and 15.5°C nighttime temperature, 15-hr day, and 500 μM light intensity. Tomato plants were grown in a greenhouse under 28°C daytime temperature, 19°C nighttime temperature, 15-hr days, and 600 μM light intensity. All plants were grown at University of Georgia, Athens, GA, USA.

## Author contributions

CL and CRB conceived the study. CL and JCW performed initial optimizations. All authors performed the experiments. CL prepared figures and drafted the manuscript with input from all authors.

## Acknowledgement

The authors would like to thank Dr. Alexandre P. Marand for advice regarding optimization of the protocol. The authors also thank Julie Nelson for assistance with flow cytometry. This work is funded by the Georgia Research Alliance and the University of Georgia (CRB).

## References

“10x Genomics Chromium Single Cell Multiome ATAC + Gene Expression Kit.” https://www.10xgenomics.com/products/single-cell-multiome-atac-plus-gene-expression.

Barone, Amalia, Maria Luisa Chiusano, Maria Raffaella Ercolano, Giovanni Giuliano, Silvana Grandillo, and Luigi Frusciante. 2008. “Structural and Functional Genomics of Tomato.” International Journal of Plant Genomics 2008 (January): 1–12. https://doi.org/10.1155/2008/820274.

Buenrostro, Jason D, Beijing Wu, Ulrike M Litzenburger, Dave Ruff, Michael L Gonzales, Michael P Snyder, Howard Y Chang, and William J Greenleaf. 2015. “Single-Cell Chromatin Accessibility Reveals Principles of Regulatory Variation.” Nature 523 (7561): 486–90. https://doi.org/10.1038/nature14590.

Castro, Mariana, and Silvia Castro. 2012. “Genome Size Variation and Incidence of Polyploidy in Scrophulariaceae Sensu Lato from the Iberian Peninsula.” AoB PLANTS, 14.

Deal, Roger B., and Steven Henikoff. 2010. “A Simple Method for Gene Expression and Chromatin Profiling of Individual Cell Types within a Tissue.” Developmental Cell 18 (6): 1030–40. https://doi.org/10.1016/j.devcel.2010.05.013.

Grandi, Fiorella C., Hailey Modi, Lucas Kampman, and M. Ryan Corces. 2022. “Chromatin Accessibility Profiling by ATAC-Seq.” Nature Protocols 17 (6): 1518–52. https://doi.org/10.1038/s41596-022-00692-9.

Grindberg, Rashel V., Joyclyn L. Yee-Greenbaum, Michael J. McConnell, Mark Novotny, Andy L. O’Shaughnessy, Georgina M. Lambert, Marcos J. Araúzo-Bravo, et al. 2013. “RNA-Sequencing from Single Nuclei.” Proceedings of the National Academy of Sciences 110 (49): 19802–7. https://doi.org/10.1073/pnas.1319700110.

Guimaraes, Guilherme, Luísa Cardoso, Helena Oliveira, Conceição Santos, Patrícia Duarte, and Mariana Sottomayor. 2012. “Cytogenetic Characterization and Genome Size of the Medicinal Plant Catharanthus Roseus (L.) G. Don.” AoB PLANTS 2012 (January). https://doi.org/10.1093/aobpla/pls002.

Haberer, Georg, Sarah Young, Arvind K Bharti, Heidrun Gundlach, Christina Raymond, Galina Fuks, Ed Butler, Klaus F X Mayer, and Joachim Messing. 2005. “Structure and Architecture of the Maize Genome1[W]” 139: 13.

Hoopes, Genevieve, Xiaoxi Meng, John P. Hamilton, Sai Reddy Achakkagari, Fernanda de Alves Freitas Guesdes, Marie E. Bolger, Joseph J. Coombs, et al. 2022. “Phased, Chromosome-Scale Genome Assemblies of Tetraploid Potato Reveal a Complex Genome, Transcriptome, and Predicted Proteome Landscape Underpinning Genetic Diversity.” Molecular Plant 15 (3): 520–36. https://doi.org/10.1016/j.molp.2022.01.003.

Huang, Shao-shan C, and Joseph R Ecker. 2018. “Piecing Together Cis-regulatory Networks: Insights from Epigenomics Studies in Plants” 10: 20.

Kashima, Yukie, Yoshitaka Sakamoto, Keiya Kaneko, Masahide Seki, Yutaka Suzuki, and Ayako Suzuki. 2020. “Single-Cell Sequencing Techniques from Individual to Multiomics Analyses.” Experimental & Molecular Medicine 52 (9): 1419–27. https://doi.org/10.1038/s12276-020-00499-2.

Kaya-Okur, Hatice S., Steven J. Wu, Christine A. Codomo, Erica S. Pledger, Terri D. Bryson, Jorja G. Henikoff, Kami Ahmad, and Steven Henikoff. 2019. “CUT&Tag for Efficient Epigenomic Profiling of Small Samples and Single Cells.” Nature Communications 10 (1): 1930. https://doi.org/10.1038/s41467-019-09982-5.

Kodiha, Mohamed, Piotr Bański, and Ursula Stochaj. 2011. “Computer-Based Fluorescence Quantification: A Novel Approach to Study Nucleolar Biology.” BMC Cell Biology 12 (1): 25. https://doi.org/10.1186/1471-2121-12-25.

Luo, Cheng, Alisdair R. Fernie, and Jianbing Yan. 2020. “Single-Cell Genomics and Epigenomics: Technologies and Applications in Plants.” Trends in Plant Science 25 (10): 1030–40. https://doi.org/10.1016/j.tplants.2020.04.016.

Marand, Alexandre P., Xuan Zhang, Julie Nelson, Pedro Augusto Braga dos Reis, and Robert J. Schmitz. 2021. “Profiling Single-Cell Chromatin Accessibility in Plants.” STAR Protocols 2 (3): 100737. https://doi.org/10.1016/j.xpro.2021.100737.

Meers, Michael P, Terri D Bryson, Jorja G Henikoff, and Steven Henikoff. 2019. “Improved CUT&RUN Chromatin Profiling Tools.” ELife 8 (June): e46314. https://doi.org/10.7554/eLife.46314.

Ouyang, Weizhi, Shiping Luan, Xu Xiang, Minrong Guo, Yan Zhang, Guoliang Li, and Xingwang Li. 2022. “Profiling Plant Histone Modification at Single-cell Resolution Using SnCUT&Tag” Plant Biotechnology Journal 20 (3): 420–22. https://doi.org/10.1111/pbi.13768.

Preissl, Sebastian, Kyle J. Gaulton, and Bing Ren. 2022. “Characterizing Cis-Regulatory Elements Using Single-Cell Epigenomics.” Nature Reviews Genetics, July. https://doi.org/10.1038/s41576-022-00509-1.

Probst, Aline V., Paul F. Fransz, Jerzy Paszkowski, and Ortrun Mittelsten Scheid. 2003. “Two Means of Transcriptional Reactivation within Heterochromatin: *Transcriptional Reactivation within Heterochromatin*.” The Plant Journal 33 (4): 743–49. https://doi.org/10.1046/j.1365-313X.2003.01667.x.

Shema, Efrat, Bradley E. Bernstein, and Jason D. Buenrostro. 2019. “Single-Cell and Single-Molecule Epigenomics to Uncover Genome Regulation at Unprecedented Resolution.” Nature Genetics 51 (1): 19–25. https://doi.org/10.1038/s41588-018-0290-x.

Shultz, J. L. 2006. “The Soybean Genome Database (SoyGD): A Browser for Display of Duplicated, Polyploid, Regions and Sequence Tagged Sites on the Integrated Physical and Genetic Maps of Glycine Max.” Nucleic Acids Research 34 (90001): D758–65. https://doi.org/10.1093/nar/gkj050.

Sikorskaite, Sidona, Minna-Liisa Rajamäki, Danas Baniulis, Vidmantas Stanys, and Jari PT Valkonen. 2013. “Protocol: Optimised Methodology for Isolation of Nuclei from Leaves of Species in the Solanaceae and Rosaceae Families.” Plant Methods 9 (1): 31. https://doi.org/10.1186/1746-4811-9-31.

Sugimoto-Shirasu, Keiko, and Keith Roberts. 2003. “‘Big It up’: Endoreduplication and Cell-Size Control in Plants.” Current Opinion in Plant Biology 6 (6): 544–53. https://doi.org/10.1016/j.pbi.2003.09.009.

Thibivilliers, Sandra, Dirk Anderson, and Marc Libault. 2020. “Isolation of Plant Root Nuclei for Single Cell RNA Sequencing.” Current Protocols in Plant Biology 5 (4). https://doi.org/10.1002/cppb.20120.

Zimin, Aleksey V, Daniela Puiu, Richard Hall, Sarah Kingan, Bernardo J Clavijo, and Steven L Salzberg. 2017. “The First Near-Complete Assembly of the Hexaploid Bread Wheat Genome, Triticum Aestivum” GigaScience 6 (11). https://doi.org/10.1093/gigascience/gix097.

